# Effective design and inference for cell sorting and sequencing based massively parallel reporter assays

**DOI:** 10.1101/2022.11.07.515414

**Authors:** Pierre-Aurélien Gilliot, Thomas E. Gorochowski

## Abstract

The ability to measure the phenotype of millions of different genetic designs using Massively Parallel Reporter Assays (MPRAs) has revolutionised our understanding of genotype-to-phenotype relationships and opened avenues for data-centric approaches to biological design. However, our knowledge of how best to design these costly experiments and the effect that our choices have on the quality of the data produced is lacking. Here, we tackle this issue by developing FORECAST, a Python package that supports the accurate simulation of cell-sorting and sequencing based MPRAs and robust maximum likelihood based inference of genetic design function from MPRA data. We use FORECAST’s capabilities to reveal rules for MPRA experimental design that help ensure accurate genotype-to-phenotype links and show how the simulation of MPRA experiments can help us better understand the limits of prediction accuracy when this data is used for training deep learning based classifiers. As the scale and scope of MPRAs grows, tools like FORECAST will help ensure we make informed decisions during their development and the most of the data produced.

## Introduction

The falling costs of DNA synthesis and sequencing are enabling new approaches to understand how biological function is genetically encoded [1–3]. Many of these emerging techniques exploit our newly found ability to create large libraries of precisely designed DNA sequences (i.e., a diverse set of genotypes) and efficiently assess how each of these sequences is linked to a cellular behaviour (i.e., their individual phenotypes) [4–7]. Unravelling such genotype-to-phenotype maps (GP maps) via data-centric approaches has lead to the creation of statistical models that are able to effectively design *de novo* genetic regulatory elements [8–11] and better understand the principles of evolution [12, 13].

Massively Parallel Reporter Assays (MPRAs) are a key development that has made this approach possible. This method involves generating a library of diverse genotypes whose phenotypes are linked to the expression of a fluorescence-based reporter. Using highly parallel measurement techniques, millions of genotype-to-phenotype relationships can then be established during a single experiment. Crucially, the scale of these assays ensures sufficient sampling of key areas in sequence space such that statistical models can capture or learn important features of the GP map.

One of the most popular MPRAs is based on Fluorescence Activated Cell Sorting (FACS) followed by deep sequencing and is commonly referred to as Flow-seq or Sort-seq (**Figure 1a**) [5, 8, 10, 12, 14–19, 19–23]. In a Flow-seq experiment, cells are engineered to link the phenotype of interest with the expression level of a fluorescent reporter protein. This pool of cells is then sorted into bins covering different levels of fluorescence, and the contents of each bin separately sequenced to determine the genotypes associated with that observed phenotype (i.e., fluorescence intensity). The ability to generate diverse genetic libraries with approaches like highly parallel chip-based oligo synthesis, and then sort and sequence hundreds of millions of cells has enabled GP maps to be produced in unprecedented detail for many genetic parts and systems [18].

**Figure 1:**
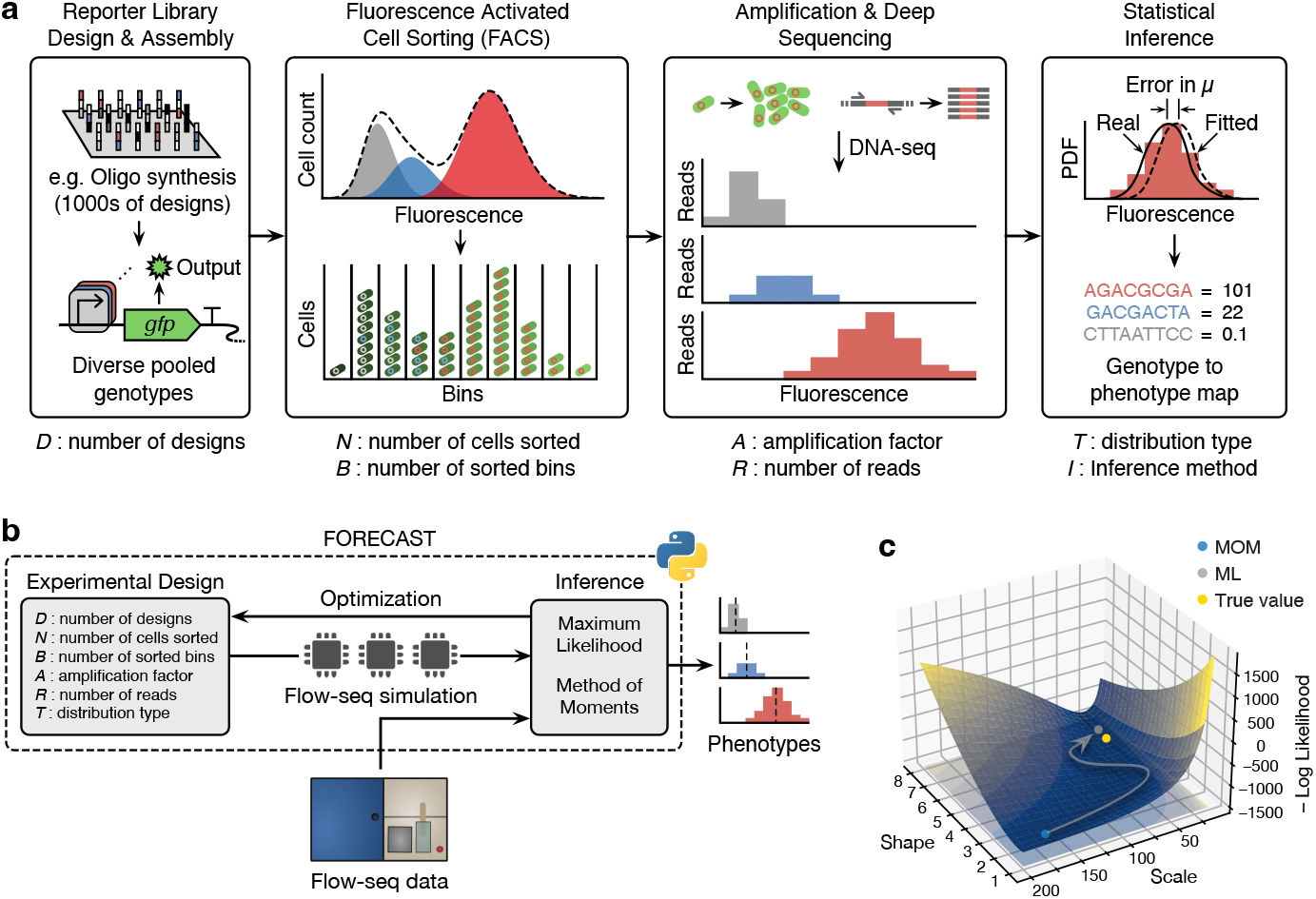
Overview of the FORECAST Python package and core functionality. (**a**) Main steps of a Flow-seq experiment. Experimental factors included in FORECAST’s probabilistic model are shown below each step. (**b**) Schematic of the FORECAST package. FORECAST allows for the definition of experimental designs that can then be simulated to generate synthetic Flowseq data. Inference of phenotype (i.e., fluorescence distribution) from real and synthetic Flow-seq data can be performed using either a Method of Moments (MOM) or Maximum Likelihood (ML) approach. (**c**) Negative log-likelihood surface computed from synthetic Flow-seq data of an example genetic variant. The ML inference method starts from the MOM estimate (blue point) and minimise the negative log-likelihood to find more accurate estimate (grey point) of the true parameter value (yellow point).

While Flow-seq based MPRAs have revolutionised our ability to map out genotype-to-phenotype landscapes, the discrete binning by cell fluorescence that is necessary during the sorting phase raises issues when trying to recover a continuous fluorescence distribution for a genetic variant and accurately quantify the uncertainty in such measurements. This is problematic, as data sets exhibiting large levels of noise are ill-suited to data-driven model identification [24]. Some experimental studies have attempted to vary key experimental factors to assess and improve measurement accuracy (e.g., varying the number of sorting bins) [14, 22, 23]. However, these have not generally been systematic in their approach. Therefore, the numerous options available when developing a new Flow-seq based MPRA are typically chosen arbitrarily or to mimic previous studies. Furthermore, the methods used to infer fluorescence distributions from discretely binned Flow-seq data are often basic and can lead to biased results. More accurate estimators have been derived, but their complexity has hindered broad uptake. [10, 25].

In this work, we aim to address these challenges by developing a Python package called FORECAST that implements an efficient Maximum Likelihood (ML) based inference method and a probabilistic model of Flow-seq experiments to allow for the systematic computational exploration of parameters associated with both the experimental and analysis procedures. We demonstrate the ability of FORECAST to uncover the most critical choices when designing a Flow-seq experiment to ensure accurate inference of fluorescence distributions, and show how we can more rigorously explain the performance and source of errors for deep learning based models trained using real Flowseq data sets. The ability of FORECAST to both aid in the design of new Flow-seq experiments and accurately analyse existing data sets makes it a valuable tool for supporting the creation of accurate models to both engineer and understand the genetic encoding of biological functions.

## Results

### Overview of FORECAST

The FORECAST package implements two core functionalities (**Figure 1b**). The first is a probabilistic simulator of Flow-seq experiments, where all major parameters for an experimental design can be varied. Each step in a Flow-seq experiment is considered covering: the diversity of the genetic library that acts as input (i.e., the number of unique genetic variants), the sampling and sorting of a mixed pool of cells harbouring this library, the amplification of sorted designs by cell growth or PCR, and the final sequencing step that captures for each genetic variant the number of reads recovered from each bin (**Methods**).

In addition to the simulation of experiments, FORECAST also implements several inference methods for estimating precise fluorescence distributions from discretely binned Flow-seq data. We provide two options for inference that use either a Method of Moments (MOM) or Maximum Likelihood (ML) based approach (**Figure 1c**). The MOM approach is mostly agnostic to the data-generating process, while the ML approach explicitly models the Flow-seq sorting and sequencing steps in its likelihood function. Practically, the MOM approach finds estimates by a weighted average of the binned data, whereas the ML approach minimises the likelihood function of an inhomogeneous Poisson process. Both estimators allow for fluorescence of a particular genetic variant to take the form of either a Gamma or Log-normal distribution to capture the inherent variation that is present in gene expression across a population of cells [26–28] (**Methods**). These inference methods can be easily applied to existing Flow-seq data sets or FORECAST-generated computational simulations.

### Improved inference for Flow-seq data

A critical step in a Flow-seq experiment is the inference of a continuous fluorescence distribution from the discretely binned sequencing data. It is common for this step to make use of a simple MOM estimator where a mid-point fluorescence value for each sorting bin is weighted according to the relative read frequencies across them. Although this approach is known to be unreliable and susceptible to noise in fluorescence measurements, it is still a common method due to the ease of implementation [18, 25, 29]. An alternative and more robust method is to use an ML based approach [10, 25], but efficient implementation of this method can be difficult and the benefits compared to MOM in this context are not well understood.

To better understand the impact of the inference method on the quality of the recovered distributions from Flow-seq experiments, we made use of the simulation capabilities of FORECAST to generate a large and biologically realistic Flow-seq data set with a known ground truth for every genetic variant in the library. We specifically simulated a Flow-seq experiment with a library containing 1018 genetic variants that had Gamma distributed fluorescence, mimicking biologically realistic protein expression characteristics [26] (**Methods**). In this experiment, we sorted 10^6^ cells into 8 bins, amplified the sorted cells by a factor of 1000, and performed sequencing using 10^7^ reads. We then conducted inference on this synthetic Flow-seq data using both the MOM and ML estimators implemented in FORECAST.

The accuracy of both inference methods was assessed by comparing the mean and standard deviation (SD) of the estimated fluorescence distribution for each genetic variant with the known ground truth (**Figure 2a,b**). We found that the ML estimates closely matched the ground truth with a Mean Absolute Percentage Error (MAPE) of 3% and 5% for the mean and SD, respectively. In contrast, the MOM estimates were continually higher than the ground truth with MAPE values of 42% and 81% for the mean and SD, respectively. Furthermore, the MOM estimates displayed a ‘staircase’ pattern that was highly pronounced for the SD estimates (**Figure 2b**). More detailed analysis of this pattern revealed that MOM estimates of fluorescence SD are similar for genetic variants sorted in the same bins (**Figure 2b,f,g**), but vary strongly when cells fall across multiple bins (**Figure 2b,h**). This artifact results from the fact that the MOM method assumes a fixed (centre point) value of fluorescence for every design in a bin, which makes it highly sensitive to cells for a design falling near bin boundaries. We confirmed that binning was the cause of this feature by running additional simulations with an increasing number of bins. As expected, this resulted in a larger number of ‘stairs’ in the inferred mean and SD values across the library of designs (**Supplementary Figure 1**).

**Figure 2:**
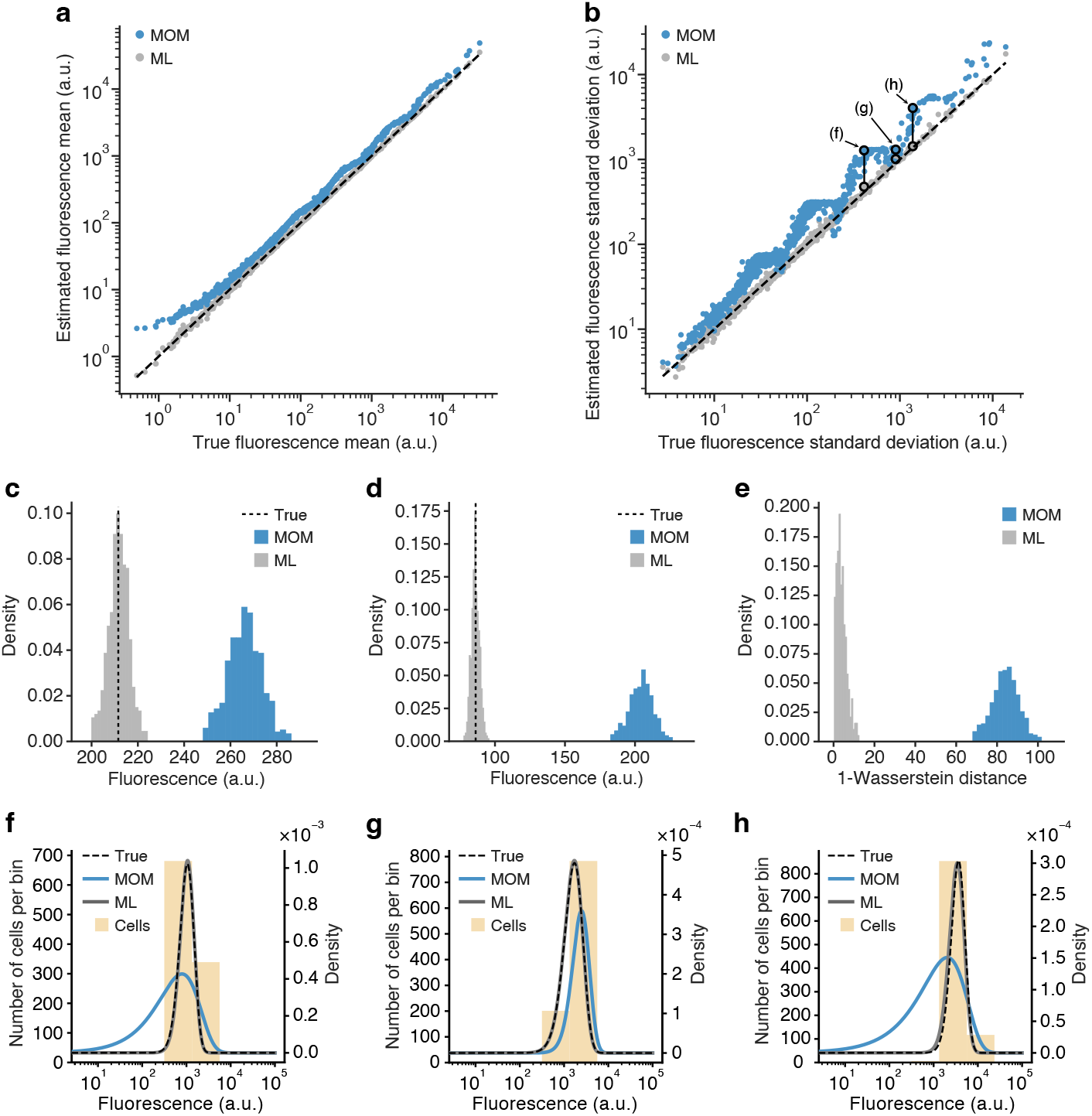
Characterizing the performance of the inference methods. (**a**) Scatter plot of estimated versus true mean fluorescence values from a simulated Flow-seq experiment of 1018 genetic variants. Each filled point corresponds to a genetic variant, blue for MOM estimates and grey for ML estimates. Black dashed lines denotes *x* = *y* (i.e., perfect accuracy). (**b**) Scatter plot of estimated versus true standard deviation (SD) in fluorescence for the same Flow-seq simulation. Each filled point corresponds to a genetic variant, blue for MOM estimates and grey for ML estimates. Black dashed lines denotes *x* = *y* (i.e., perfect accuracy). Pairs of annotated points highlighted by an unfilled black circle and connected by a line correspond to the MOM (top) and ML (bottom) estimates for the same genetic variant. Annotation corresponds to panel displaying the underlying distributions (**f–h**). (**c**) Sampled distribution of the fluorescence mean inferred using the MOM (blue) and ML (grey) approach for 500 separate Flow-seq experiments with the true value shown as a black dashed line for the ‘mob’ genetic variant. (**d**) Sampled distribution of the fluorescence SD inferred using the MOM (blue) and ML (grey) approach for the same replicate Flow-seq experiments with the true value shown as a black dashed line for the ‘mob’ genetic variant. (**e**) Sampled distribution of the 1-Wasserstein distance inferred using the MOM (blue) and ML (grey) approach for the same replicate Flow-seq experiments for the ‘mob’ genetic variant. (**f**, **g**, **e**) Inferred (blue line for MOM and grey line for ML) and true (black dashed line) density distributions plotted on top of the discrete histogram of cells sorted (light orange) for the highlighted genetic variants in panel b.

To further characterize the statistical properties of the estimators, we measured the bias (i.e., the closeness between the estimates and the true value) and variance (i.e., the dispersion from the mean prediction) of each estimator by simulating a further 500 Flow-seq experiments for the same library and identical experimental parameters (**Methods**). The sampled distributions for the fluorescence mean and SD estimators across these replicates consistently showed smaller bias when using the ML inference method. The mean ratio (ML/MOM) for the bias was found to be 0.005 for the mean and 0.016 for the SD. An example of the sampled variation for a specific genetic variant is shown in **Figure 2c,d** and for the entire genetic library in **Supplementary Figure 2 a,b**. The variance of both estimators was comparable between the methods, with a mean ratio (ML/MOM) of 0.86 and 1.17 for the fluorescence mean and SD, respectively (**Figure 2c,d**; **Supplementary Figure 2**).

We further investigated the ability of each inference method to globally recover the underlying fluorescence distributions by measuring the 1-Wasserstein distance (WD) between the inferred and ground truth distributions. Intuitively, the WD measures the minimum cost of transforming one distribution into another and indicates the closeness between them. Using this distance metric, ML-inferred fluorescence distributions were found to always be closer to the ground truth than MOM-inferred versions, with a maximum ratio (WD_ML_/WD_MOM_) of 0.34 between the two methods. An example of the WD sampled distribution for one genetic construct is shown in **Figure 2e** and for the entire genetic library in **Supplementary Figure 2**.

An additional benefit of using the ML approach is that it provides uncertainty information in the form of confidence intervals (**Methods**). We validated the theoretical derivation of the confidence intervals by using the replicate Flow-seq simulations (**Supplementary Figure 4**) and measured the coverage rate (i.e., the number of times the confidence interval contains the true value) (**Methods**; **Supplementary Figure 5**). We found that the simulation results reliably matched the confidence intervals produced from the limited number of cells and sequencing reads used.

While these results demonstrated that ML inference provides a more accurate estimation of a genetic construct’s fluorescence distribution, it comes at a computational cost. Comparing the run-time for MOM and ML estimators revealed that our ML estimator was approximately 4,000 times slower to compute. On average for a typical desktop computer, MOM inference took ~ 77 × 10^−3^ ms to compute for a simple construct compared to ~ 340 ms when using the ML method (**Methods**). To overcome potential run-time difficulties when millions of ML estimates are required, we implemented multiprocessing in FORECAST to allow for the ML computations to be parallelized. We tested the scaling characteristics of our code from 1 to 28 processor cores and found a linear scaling (factor = 0.6) with no decrease in the ability to utilize newly available cores (**Supplementary Figure 6**).

### Optimal design of Flow-seq experiments

Flow-seq experiments are costly and time-consuming to perform. Yet, their design is often arbitrarily based on other studies, which provides no guarantee as to the quality or accuracy of the data that will be produced. This stems from difficulties in testing the effect that experimental parameters will have on the ability to infer accurate phenotypes from data. To address this, we again employed FORECAST’s ability to simulate Flow-seq experiments and systematically explored the experimental design space to better understand the influence of each experimental factor (**Methods**). We varied the number of cells sorted and sequencing reads per genetic variant, as well as the number of logarithmically spaced bins used for cell sorting, as these are the key factors that map to common choices when carrying out an Flow-seq experiment (e.g., the types of sequencer used or setting on a FACS machine). We also used both the MOM and ML estimators to infer the fluorescence distribution for each experimental design and characterized the error between the inferred and ground truth for every genetic variant as a measure of accuracy.

We first explored the impact of varying each experimental parameter on the global accuracy of the two inference methods by computing the mean of the WD (MWD) across the entire genetic variant library. Increasing the number of cells sorted, sequencing reads, and sorting bins all improved the accuracy of the inferred values (i.e., lower MWD values; **Figure 3a-d**). Furthermore, inference using ML consistently outperformed the MOM approach, with the gap in performance becoming reduced as the number of sorting bins increased. We also see a reduction in MWD for the ML estimates as the number of reads and cells per genetic variant increases. This is coherent with the asymptotic consistency of ML estimates, supporting the fact that this method can recover an underlying distribution with arbitrary precision [30].

**Figure 3:**
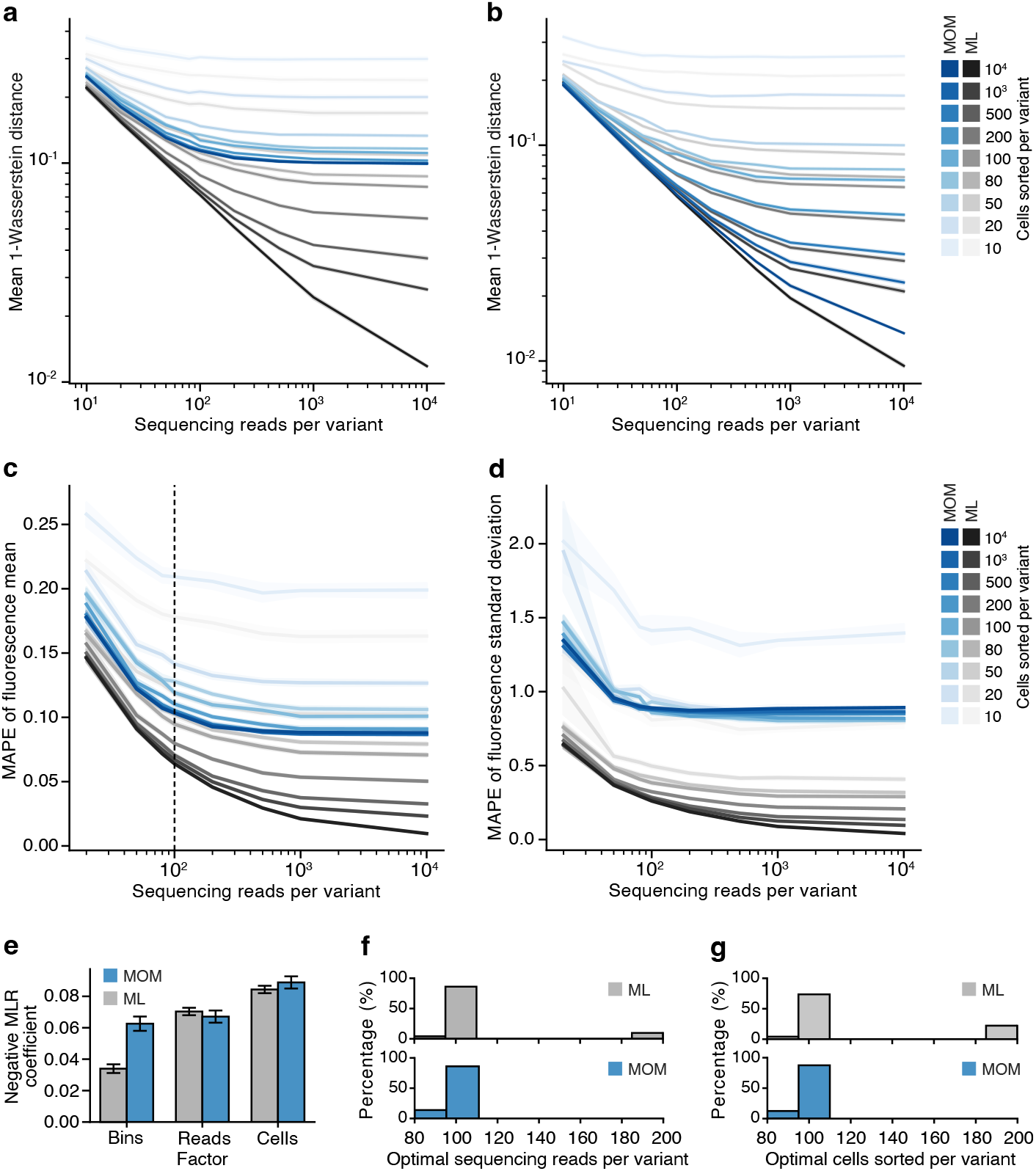
Effect of experimental design parameters on the accuracy of phenotype inference for Flow-seq experiments. (**a**) Global accuracy given by the Mean 1-Wasserstein distance (MWD) of a simulated Flow-seq experiments with 8 bins used for cell sorting and varying numbers of sequencing reads per genetic variant and numbers of cells sorted per variant. MWD shown for inference using Method of Moments (MOM; blue lines) and Maximum Likelihood (ML; grey lines) and standard deviation in these measurements shown by shared regions around each line. (**b**) Global accuracy in terms of MWD shown for same experiments as in panel a, but using 32 bins for cell sorting. (**c**) Mean Absolute Percentage Error (MAPE) when estimating the fluorescence mean using either MOM (blue) or ML (grey) inference methods. Averages were taken over the full library of genetic variants and Flow-seq simulation replicates. Black dashed line denotes the general optimal operating point. (**d**) MAPE when estimating the fluorescence standard deviation (SD) using either MOM (blue) or ML (grey) inference methods. Averages were taken over the full library of genetic variants and Flow-seq simulation replicates. Coefficients of the multiple linear regression (MLR) used to fit the MWD as a function of the number of sorting bins (‘Bins’), the log-number of reads (‘Reads’) and log-number of cells per genetic variant (‘Cells’). (f) Distribution of the optimal operating point in terms of the number of sequencing reads per genetic variant in a Flow-seq experiment for MOM (blue, bottom) and ML (grey, top) inference methods. (g) Distribution of the optimal operating point in terms of the number of cells sorted per genetic variant in a Flow-seq experiment for MOM (blue, bottom) and ML (grey, top) inference methods.

Similar observations are seen for the local behavior of each inference method, which was assessed using the Mean Absolute Percentage Error (MAPE) of the fluorescence mean or SD (**Figure 3c,d**). We found that the relative error made using the ML approach is always smaller compared with the MOM approach (**Supplementary Figure 7**). Furthermore, we observed that estimating the fluorescence SD is more challenging than estimating the mean with the MAPE of the SD being 5 times larger than the MAPE of the mean for both inference methods (MAPE ratios across all sequencing reads and inference methods are provided in **Supplementary Figure 7**).

To quantify the relative importance of each experimental factor, we used a multiple linear regression model to capture the relationship between the MWD and the number of bins, sequencing reads, and cells sorted per genetic variant (**Methods**). By extracting the coefficient for each predictor variable, we could compare the importance of every experimental factor for each inference method. We found that the number of bins was twice as important for accurate MOM inference compared to when ML was used. Furthermore, increases in the number of sequencing reads and cells per genetic variant improved the accuracy of inference for both methods, although with diminishing returns (**Figure 3a**).

Our ability to observe how various experimental factors affect the accuracy of inference, offers the chance to provide rules that could help ensure a good accuracy of measurements when designing an MPRA experiment. To determine these ideal parameters, we analyzed the diminishing returns phenomena observed when increasing each experimental parameter (**Figure 3a,b**). We found that a constant decrease in MWD required increasing resources. Therefore, we would expect a point where the relative cost in terms of the scale of an experiment is no longer worth the corresponding benefit in improved measurement accuracy. By assuming constant relative costs, we determined the optimal operating points for each experimental factor using the kneedle algorithm [31, 32] (**Figure 3a**; **Methods**). It should be noted that this approach could be extended to account for varying costs of factors in the experimental design by weighting each of these accordingly.

To extract the optimal number of reads, we plotted the MWD as a function of the normalized number of reads and located the optimal operating point (**Supplementary Figure 8**). The MWD was taken by averaging over all simulations from the factorial experiment (**Methods**). Similarly, we extracted the optimal number of cells by plotting MWD as a function of the normalized number of cells and using the same procedure (**Supplementary Figure 8**). Finally, we found the optimal number of bins by plotting the MWD as a function of the number of bins, while setting the number of cells and reads to their optimal value. This process was repeated across all simulated data from the factorial experiment, and we found that the most frequent operating point was 100 for both the number of cells and sequencing reads per genetic variant (**Figure 3f,g**). This combination of parameters corresponds to sequencing on average all cells in the library before amplification. Using this operating point for the sequencing reads and cells, we found that 10 bins was optimal for cell sorting (**Methods**; **Supplementary Figure 9**). While these general rules are useful for guiding future experiments, the response curves we have generated can also be used to select experimental parameters where specific costs and trade-offs exist for a particular experimental setup and a known level of accuracy is required.

### Unraveling sources of errors when using Flow-seq data

The large amounts of data produced by Flow-seq experiments are ideal for training deep learning-based models [10, 22, 33]. A challenge in this context is knowing what sort of accuracy can be achieved by a model given the inherent variability in the underlying biology and experimental procedures.

To show how FORECAST can help to answer this, we considered a recent study by Evfratov *et al*. that investigated how a messenger RNA’s 5’ untranslated region (5’UTR) regulates the translation initiation rate of an associated reporter gene [16]. This study involved a Flow-seq experiment in which a library of ~ 28, 000 different genetic variants was tested. Using data from this experiment as our ground truth, we defined a classification task where the bin (1 to 8) in which the majority of cells were sorted (i.e., the ‘mode bin’) needed to be predicted for each genetic variant. This task and the amount of available data are well suited to a deep learning-based classifier and provided an opportunity to systematically assess how experimental design choices might limit the predictive accuracy that could be achieved and pinpoint the sources of errors. Due to the stochastic nature of the sorting and sequencing steps in a Flowseq experiment, we would expect some mode bin measurements to potentially vary between experiments (especially for genetic variants whose mode fluorescence falls near bin boundaries). To estimate the experimental-level accuracy that can be achieved due to this effect, we simulated many Flow-seq experiments with a varying number of sequencing reads and sorted cells for each genetic variant (**Methods**). For each genetic variant and combination of experimental factors, we measured the mode bin and compared it to the ground truth. We then obtained a frequency of correct mode bin measurements for each experimental factor by averaging over the genetic library and simulation replicates. We found that measuring the mode bin from Flow-seq data can be difficult, especially in regimes of low-experimental resources where the frequency of correct mode bin measurements can be as low as 50% (**Figure 4a**; **Methods**). Increasing the number of sequencing reads and cells sorted was found to improve the accuracy of mode bin measurements, although with diminishing returns.

**Figure 4:**
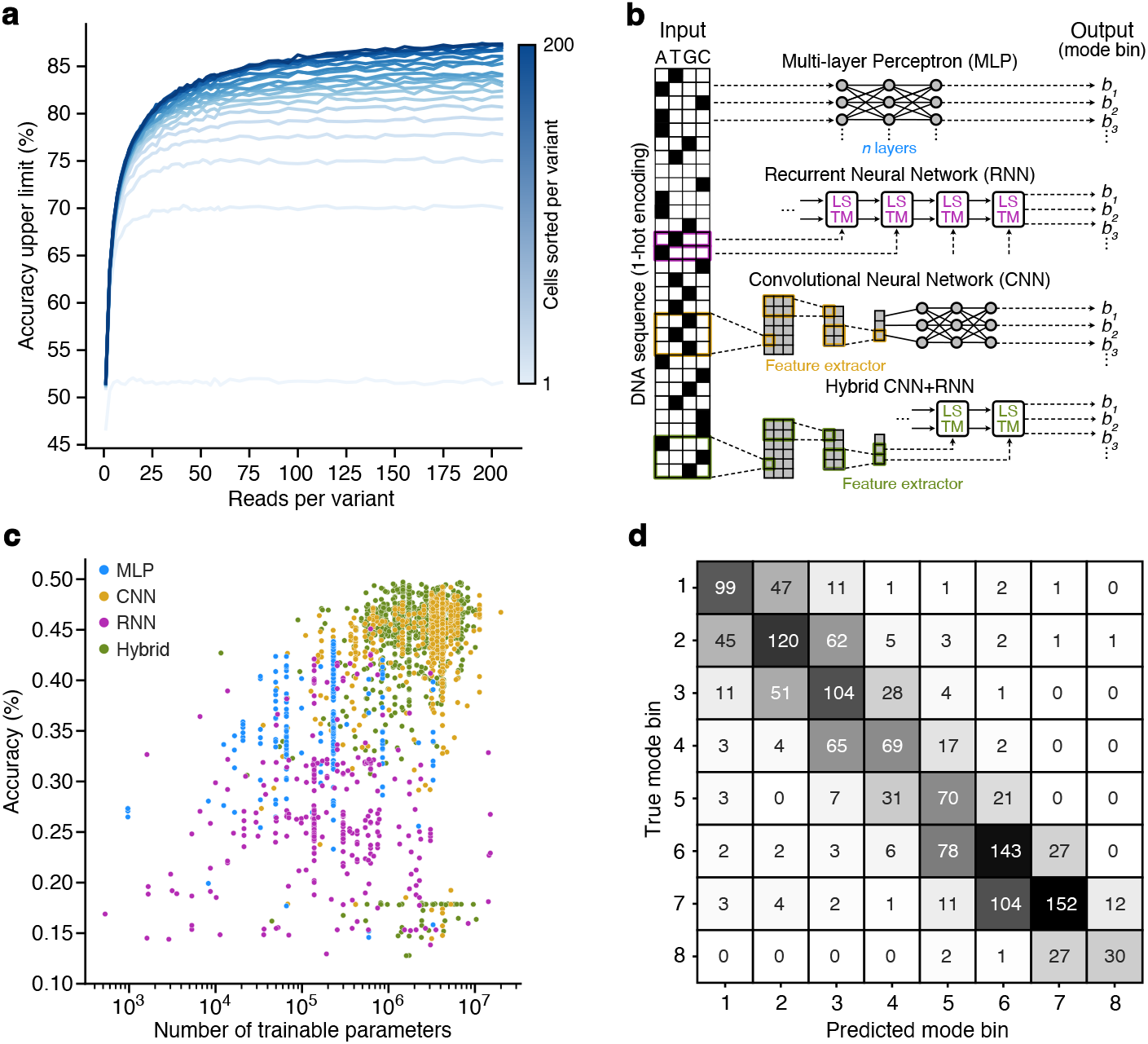
Understanding the limits to prediction accuracy from Flow-seq experiments. (**a**) Accuracy upper limit when predicting the mode bin as a function of the number of sequencing reads and cells sorted per genetic variant. Cells sorted per variant took the values 1, 6, 11, 16, 21, 26, 31, 36, 41, 46, 51, 56, 72, 80, 100, 120, 140, 160, 180, 200 (light to dark blue lines). Each value was estimated by running FORECAST simulations with different experimental factors. (**b**) Neural network architectures used to predict the mode bin from 5’ untranslated region (5’UTR) sequence. **(c)** Scatter plot of the validation set accuracy against the number of trainable parameters in the model. Each point corresponds to a particular combination of hyperparameters for a class of neural network (MLP = blue; CNN = orange; RNN = purple, Hybrid = green). (d) Confusion matrix of the hybrid CNN+RNN architecture with the best performance and fewest trained parameters evaluated on a previous Flow-seq study [16]. Global accuracy is 51.8%.

These estimates for the frequency of correct mode bin assignments allow us to quantify the maximum accuracy that can be reached for any model trained or fitted to the data from this Flow-seq study. To empirically assess this, we split the data into a training set (80%), validation set (10 %), test set (10%), and then computed the maximum achievable accuracy of a model on each data split by pulling the corresponding frequency of correct mode bin measurements given the experimental resources it was calculated from. To ensure some differences in the achievable accuracies, the test set was selected to have a higher maximum accuracy (75% compared to 70% for both the training and validation sets) by selecting genetic variants with the most sequencing reads (**Methods**).

Having selected appropriate data sets, we next needed a model to learn how 5’UTR sequence mapped to protein expression level. While mechanistic models of translation initiation do exist [34, 35], they typically are not highly predictive when applied to data sets not used for fitting. In contrast, machine-learning based models using deep neural networks have made large strides in this domain, allowing for automatic feature extraction and the best predictive accuracies seen to date [36–38].

Here, we selected four common deep neural network architectures to learn the mapping between 5’UTR sequence and the translation efficiency (i.e. fluorescence) from data. In particular, we considered a Multi-Layer Perceptron (MLP), Convolutional Neural Network (CNN) using an MLP for feature expraction as a final step, Recurrent Neural Network (RNN) (**Figure 4b**) and hybrid CNN+RNN. All of these neural network architectures were trained to predict the mode bin from the 5’UTR sequence on the training set, and a medium-sized hyperparameter optimization on the validation set was used to compare the performance of each architecture (**Methods**). We found that the best-performing architecture was the hybrid CNN+RNN, with a validation accuracy of 48%, compared to a possible maximum accuracy of 70% (**Figure 4c**).

At least two reasons could explain the 22% gap between the validation accuracy and the maximum achievable accuracy. Firstly, the approximations used to model Flow-seq experiments might be too simplistic, meaning the estimates for the maximum achievable accuracy are overly optimistic. Secondly, the neural network model might need further refinement and additional training data to better approximate the mapping between sequence and function.

To explore the first hypothesis, we evaluated the performance of the hybrid CNN+RNN model that had minimal model complexity (i.e., number of trainable parameters) while maximizing validation accuracy on the test data set (**Figure 4c**). We found that this architecture achieves a ~4% improvement in accuracy on the test set (51.8% accuracy on the test set compared to 48.3% on the validation set; **Supplementary Figure 10**), which closely matches the 5% increase in accuracy predicted by the simulations (maximum accuracy limit of 75% on the test set compared to 70% on the validations set). This suggests that the magnitude of the experimental-level accuracy estimates are likely correct and that further data is required for the model to better capture the complex relationship between sequence and function.

In addition, by plotting the confusion matrix, we observed that the distribution of errors was not random, with most errors corresponding to predictions ±1 bin away from the true mode bin (**Figure 4d**). This type of error is coherent with our simulations, as inaccurate mode bin measurements are typically in a neighboring bin of the true mode bin. In summary, the ability to estimate limits on achievable accuracies by simulating actual Flow-seq experiments provides valuable insight into what we can expect to achieve with state-of-the-art models and how model performance is best assessed.

## Discussion

As data-centric approaches to biological design and understanding rise in popularity, it is vital to make the best use of the costly and time-consuming experiments that fuel them. Here, we have shown how FORECAST can support both effective experimental design and improved inference when using large-scale datasets from Flow-seq based MPRAs. Through extensive simulation, we demonstrated how increases in the scale of a Flow-seq experiment resulted in diminishing returns for the accuracy of inferred phenotypes. This allowed us to derive some simple advice to ensure good quality data is produced from an Flow-seq based MPRA: a library with *n* different genetic variants should at least be sorted in 10 bins, with 100 cells and sequencing reads per variant. We see this recommendation as an easy-to-apply rule to help design effective Flow-seq experiments that typically sets a limit on the size of the library that can be considered given the available sorting and sequencing resources. For example, a lab using a MiSeq sequencer offering ~20 million reads per run can characterise a library containing a maximum of ~200,000 different genetic variants (assuming there is no bias in the initial library composition).

Ensuring the most is made of this data is also crucial and we have shown how the ML-based inference method implemented by FORECAST provides a 16.2-fold improvement in MAPE compared to widely used MOM based approaches. Such improvements are supported by other studies that have applied ML to inference to MPRAs, but which did not account for the stochasticity introduced by the sequencing step [25]. The use of our ML estimator is not only compelling due to its ability to reach high inference accuracy from smaller-scale experiments, but it is also able to provide valuable information on the uncertainty of the estimates in the form of confidence intervals. Furthermore, existing Flow-seq data sets can benefit from this easy to use ML inference functionality, offering the potential to refine our understanding of core biological processes like transcription [10, 14, 17, 39–42] and translation [8, 12, 17, 20, 23, 33, 43] via reanalysis.

Although FORECAST offers many benefits, there are some limitations to its use. The most prominent of these is the limited number of distributions available for describing the inherent variability in reporter expression for simulation and inference. The Log-normal and Gamma distributions included will cover most use cases. However, some biological processes can lead to complex expression profiles with bimodal features that are difficult to capture in a parametric form [8]. The open source nature of FORECAST’s code allows for other distributions to be implemented for specific use cases (**Data Availability**), and an interesting future direction would be to include goodness of fit functionality (e.g., using a chi-squared test) to fit the distribution most appropriate to the data at hand.

In summary, this work demonstrates how computational methods can support many aspects of experimental and model design, allowing for the exploration of design areas that would otherwise be impractical or too expensive to investigate physically. As the scale of MPRAs increases, making the best use of the resources available and the data generated will be critical for unraveling accurate GP maps and offering new approaches to rationally engineering biology.

## Methods

### Data-generating process

We considered the following generating process for each Flow-seq simulation. Let *D* be the diversity of the library (the number of different genetic variants), *N* the number of FACS events (i.e., the total number of sorted cells), *B* the number of bins used to sort the cells, [*F_j_*, *F*_*j*+1_] the fluorescence bounds of sorting bin *j* and *R* the total number of sequencing reads. We use the lowercase versions of each parameter to refer to any value in between. The cell sorting step is modelled by the first loop, while the second loop models the sequencing step.

### Verifying assumptions during simulation and inference

A multinomial sampling scheme (drawing with replacement) was used to model the sequencing step in the Flow-seq simulations. Although an hypergeometric sampling scheme (drawing without replacement) would be more appropriate, it would create a significant computational bottleneck during our simulations. Using a multinomial sampling scheme allows us to generate faster simulations, but we need to verify this approximation.

**Algorithm 1.**
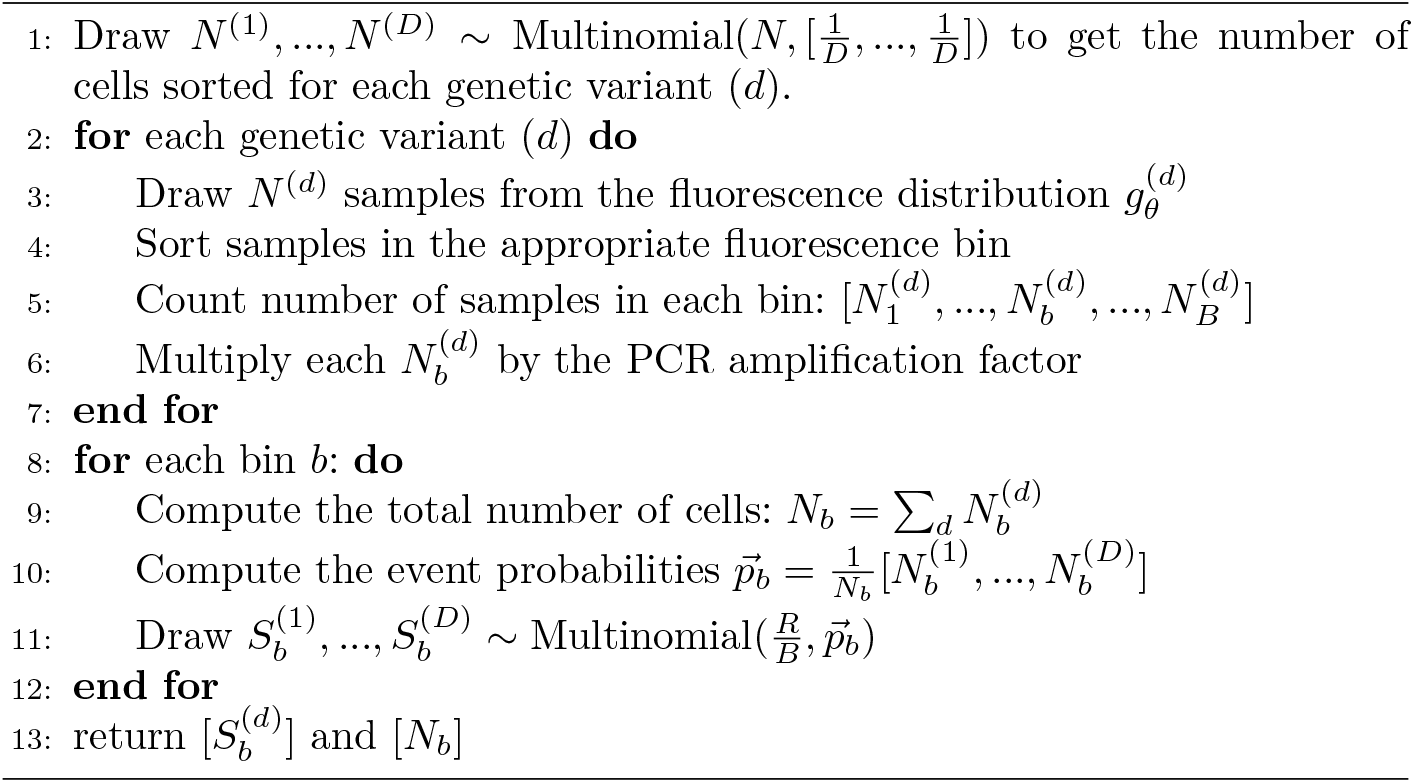
Generate Flow-Seq data

The validity of the multinomial approximation is controlled by the ratio 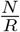, where *N* is the total number of cells sorted and *R* the total number of sequencing reads. Hence, conclusions should become particularly accurate as *N* > 10*R*. We used a second approximation to derive the log-likelihood, which is to use a Poisson distribution instead of a bimonial distribution. This approximation requires to be valid:

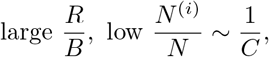

where *B* is the number of sorting bins, *C* the diversity of the library and *N*^(*i*)^ the number of cells sorted corresponding to the *j*-th genetic variant. This approximation is well founded because we typically have for common Flow-seq (and during our experiments) :

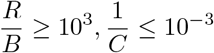

### Fluorescence data and parameterization

Here we specify the origin of the fluorescence data we use for our simulations and the families of distributions used to model the fluorescence. To ensure resemblance with reality, we used experimental Flow-seq data and *E. coli* fluorescence distribution parameters to generate our simulations. Gamma and Log-normal distributions were used separately to model the fluorescence data of each genetic variant. We chose the shape/scale parameterization when using the gamma distribution to model the fluorescence. This means that the density function *g*(*x*), the mean *μ* and the standard deviation *σ* are given by:

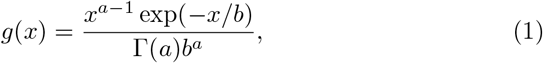

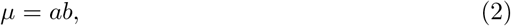

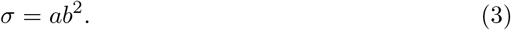

Gamma distribution parameters (*a*, *b*) for each genetic variant were taken from existing data [26]. We simplified the analysis when using the Log-normal distribution modelling choice by first log-transforming the fluorescence data. As we operate on the log-fluorescence domain, the Log-normal distribution has the same parameterization as a Gaussian distribution. The density function *g*(*x*) is then given by,

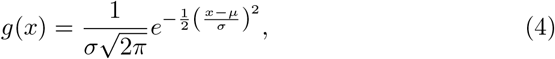

where *μ* is the mean log fluorescence and *σ* is the standard deviation of the log fluorescence. Parameters were inferred from a previous Flow-Seq experiment [8].

### Inference using the Method of Moments

During the sorting step, the precise fluorescence information for each cell is lost. In the MOM inference procedure, each cell is assigned a fluorescence value depending on the bin it was sorted in; here we used the fluorescence midpoint of each bin. This enables us to compute the experimental mean 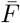 and standard deviation *σ_F_*. The MOM estimates 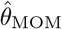 are then obtained by equating the experimental and theoretical moments:

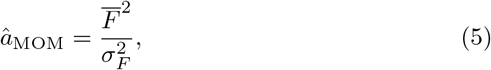

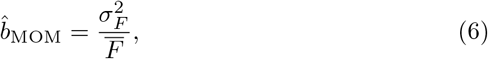

or

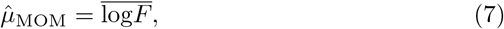

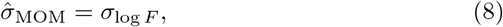

where 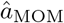 and 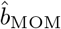 are the estimators for the shape and scale of the Gamma distribution in Equation 1 and 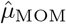 and 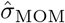 are the estimators for the parameters of the Gaussian distribution in Equation 4.

### Deriving the model likelihood

We extended a previous formulation of the likelihood used to account for the stochasticity observed during the sequencing step [10, 25]. A typical model to represent the sequencing count data is to use a Poisson distribution [44–46]. We denote the read count for genetic variant (*d*) observed when sequencing bin *b* by 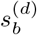. Noting that 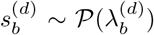, the data generating procedure can then be written as,

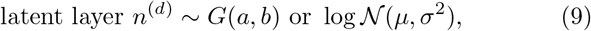

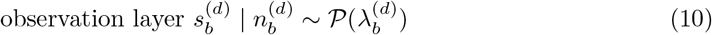

Let 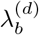 be the rate of the Poisson distribution: 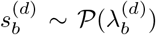. This rate is given by,

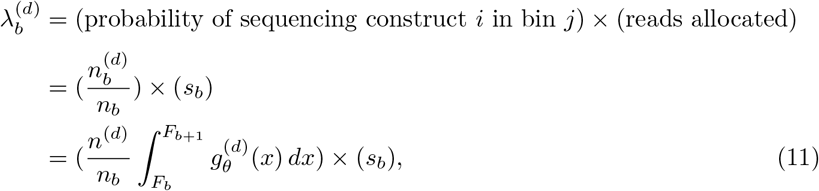

where 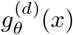 is either a gamma or lognormal density function, [*F_b_*; *F*_*b*+1_] are the fluorescence boundaries for sorting bin *b*, 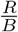 is the ratio of sequencing reads allocated per sorting bin, *n*^(*d*)^ the number of cells sorted for to the genetic variant (*d*) and *n_b_* the number of cells sorted in bin *b*. A common estimate for *n*^(*d*)^ is,

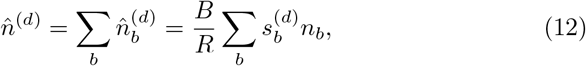

where *n_b_* indicates the number of cells sorted in bin *b* [10].

The output of a Flow-seq experiment is the sequencing data, which can be represented by a *D* × *B* matrix of discrete values. The likelihood of observing one row of this matrix is the product of the probabilities that we sequenced 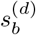 times the genetic variant (*d*) in bin *b*, thus,

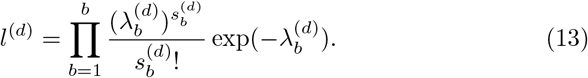

For numerical stability, we worked with the negative log-likelihood giving,

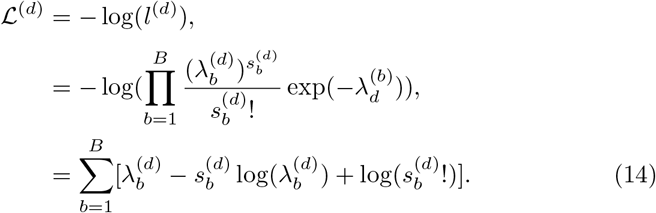

### Finding the likelihood maximum

The point *θ*_ML_ in parameter space maximizing the likelihood function 13 corresponds to the maximum likelihood estimates 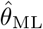. However, the first-order conditions of the likelihood function cannot be solved analytically. Therefore, we obtained the maximum likelihood estimates for the fluorescence distribution parameters by numerically minimizing the negative log-likelihood function 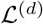 using the Nelder-Mead simplex algorithm. The starting point for the minimization procedure was initialized with the MOM estimates 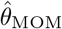. To make sure the estimates were positive after the final minimization step, we worked with the log-transformed variables log(*θ*) = (log(*μ*), log(*σ*))) when using the normal distribution and log(*θ*) = (log(*a*), log(*b*)) when using the gamma distribution. The invariance property of maximum likelihood [30] ensures the log-likelihood maximum is the same in log space. We then applied the exponential function to get the ML estimates 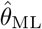.

### Computing confidence intervals

The maximum likelihood framework provides additional information on the estimates in the form of confidence intervals [30]. To obtain the confidence intervals, we computed the observed Fischer information matrix I(*θ*_ML_) by evaluating the Hessian of the negative log-likelihood 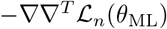 at the point maximizing the likelihood. We then computed the inverse of the observed Fischer information I(*θ_ML_*)^−1^ and extracted its diagonals elements to get the 100(1 − *α*)% confidence intervals for each estimate 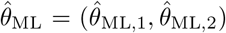. Specifically,

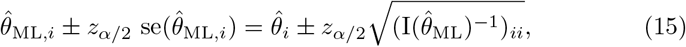

where *z*_*α*/2_ is the corresponding z-score for the 100(1−*α*)% confidence interval. To compute the confidence intervals for the mean *μ* and the variance *σ*^2^ when using the gamma distribution, we use the delta theorem and the transformation *g* : (*a, b*) → (*ab, ab*^2^). This leads to the following 100(1 − *α*)% confidence interval,

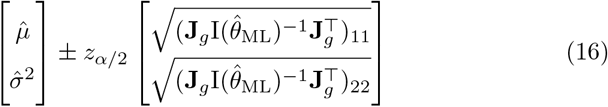

with **J**_*g*_ the Jacobian of the transformation

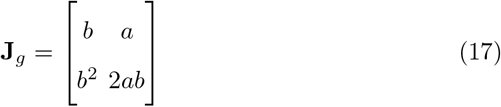

### Evaluation of estimates

After performing simulations and calculating the estimates, we considered several criteria to evaluate the performance of the inference at the genetic variant level. Assessment of global accuracy was conducted using the WD between the known ground truth and inferred distributions. WD was computed empirically by sampling 100,000 points from the inferred and ground-truth distributions. Parameter-level accuracy was assessed by measuring the percentage error (PE) given by,

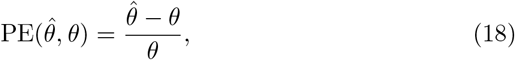

where 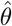 is any estimate of interest (e.g., mean fluorescence or shape parameter) and *θ* the corresponding ground truth. Library-level performance was assessed by averaging the metrics over all genetic variants to compute the MWD and MAPE given by,

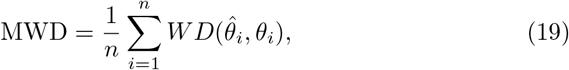

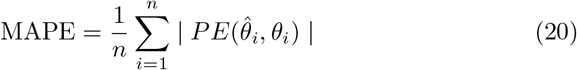

Here, *i* is the index or the genetic variant and *n* is the size of the library. Coverage of the 95%-confidence interval was assessed by computing the proportion of times the confidence interval contained the correct parameter for 500 replicates. This coverage rate was deemed acceptable when it was within two Standard Errors (SE) of the nominal coverage probability. With 500 replicates, 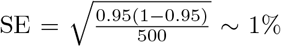 and hence an acceptable coverage has to be within [95% − 1.96 × 1%, 95% + 1.96 × 1%] ~ [93%, 97%].

### Assessing inference quality

After the inference step, it was necessary to develop a systematic procedure to inspect the reliability of estimates. Border effects, where fluorescence values were near zero or near the maximum of the final sorting bin, were detected by computing the percentage of reads located in the first and last sorting bin. Only variants with percentage less than 70% were retained. Situations where only a few reads were recovered were addressed by removing constructs with sequencing reads falling exclusively in one sorting bin. Model specification/numerical issues were identified and removed by dropping genetic variants that had non-invertible Hessians or when the confidence intervals were bigger than the corresponding estimates.

### Experimental factors

Five experimental factors were selected for evaluation of their impact on the accuracy of the statistical inference: 1. the number of different genetic variants in the library *D*, 2. the number of sorted cells *N* in the library, 3. the number of reads *R* allocated during the sequencing step, 4. the number of bins *B* used during the sorting step, and 5. the fluorescence distribution modelling choice *Z*. Normalised versions of the number of reads and the number of sorted cells were obtained by taking their ratios with the diversity of the assay. These normalised values are indicated with tildes 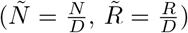 and remove the dependence of the assay diversity *D* on the conclusions.

### Pipeline to characterise inference methods

To characterize the statistical behaviour of both MOM and ML estimator in a simple setting, we performed one round of simulation+inference protocol with the following values for the experimental factors: Library of 1018 genetic variants with gamma fluorescence distributions, 10^7^ sequencing reads, 10^6^ cells sorted, 10^2^ PCR amplification factor, 8 bins and maximum fluorescence (f_max_ = 10^5^). Bias and variance of each estimator was measured by generating 499 additional synthetic Flow-seq datasets with identical experimental factors using a different random seed. We reported the sampling variation associated with the mob protein from the Taniguchi *et al*. dataset [26].

### Factorial experiment

To study both individual and interaction effects between experimental factors, we conducted a factorial experiment over a comprehensive range of experimental factors. The values we explored for the number of cells sorted and the number of sequencing reads were 10^4^, 2 × 10^4^, 5 × 10^4^, 8 × 10^4^, 10^5^, 2 × 10^5^, 5 × 10^5^, 10^6^ and 10^7^. For cell sorting, we considered 6, 8, 10, 12, 14, 16, 18, 20, 32 bins and estimated the fluorescence distribution using both Log-normal and Gamma modeling choice. Each combination of experimental factors was replicated 15 times with a different random seed to account for sampling variation, which amounted to 21,870 simulated Flow-seq experiments assessing ~22 million genetic variants.

### Comparing experimental factors

We fitted two multiple linear regression models on a subset of the factorial experiment dataset to quantify the relative importance of each experimental factor to the accuracy of the estimates. The response variable was the MWD, using either the ML or MOM inference method. The predictor variables are the number of bins, the log-transformed number of reads per genetic variant and the log-transformed number of cells per genetic variant. The data used to fit the model was restricted to values where this relationship is approximately linear to satisfy the assumptions from linear regression. Each predictor variable was min-max normalized.

### Deep learning data

Flow-Seq sequencing data was extracted from a previous study [16], where the translational initiation rate of messenger RNAs across a library of constructs with different 5’UTRs was measured. Data was aggregated from experiments ‘20N’, ‘30N’ and ‘RML eno’ and ultimately comprised of 28,402 different RNA sequences. The sequencing mode bin (between 1 and 8) was determined for each RNA sequence and served as the target output. Number of cells per sequence was estimated using Flow-Cytometer population events data for each experiment (private correspondence with the authors) and the equation 12. 80/10/10% split was used to create Train/Validation/Test sets. Sequences with the most sequencing reads (cutoff at 40) were prioritized for the test set creation to ensure high-quality data is used for model evaluation. The input RNA sequences were one-encoded into a matrix of size 4 × 30. Data augmentation was performed on the training and validation set by zero-padding 20 nucleotide long sequences. 20 nucleotide long sequences from the test set were zero-padded on the right.

### Assessing the experimental-level accuracy of the Evfratov dataset

The target output for each genetic variant in the Evfratov dataset [16] is the sorting bin with the most cells (i.e., the ‘mode bin’). We extracted the mode bin from data for each genetic variant by computing an estimate for the number of cells sorted in each bin using Equation 12 and selecting the sorting bin corresponding to the biggest number of cells sorted. However, these estimates for the mode bin are unstable due to sampling variation occurring at the sorting and sequencing step.

To assess the accuracy of each target output from the Evfratov dataset, we used as proxy the frequency of correct mode bin measurement in a Flow-Seq with similar experimental parameters. For example, the attributed accuracy for the mode bin of a genetic variant from the Evfratov dataset characterized with 8 bins, 20 sequencing reads and 30 cells corresponds to the frequency of correct mode bin measurement of a Flow-seq experiment having on average 20 sequencing reads, 30 cells per genetic variant and using 8 sorting bins. We then averaged the accuracy of each target output to get the maximum achievable accuracy on each dataset split.

The frequency of correct mode bin measurement was measured by first simulating a Flow-seq experiment, and then counting the number of times each genetic variant was attributed the true mode bin. We used the fluorescence parameters from the Taniguchi library [26] and matched the experimental factors from the Evfratov experiment: cells were sorted into 8 bins with sequencing reads = 1, 3, 5, 7, 9, 11, 13, 15, 17, 19, 21, 23, 25, 27, 29, 31, 33, 35, 37, 39, 41, 43, 45, 47, 49, 51, 53, 55, 57, 59, 60, 65, 70, 75, 80, 85, 90, 95, 100, 105, 110, 115, 120, 125, 130, 135, 140, 145, 150, 155, 160, 165, 170, 175, 180, 185, 190, 195, 200, 205, 500 and 1000, and numbers of cells sorted = 1, 6, 11, 16, 21, 26, 31, 36, 41, 46, 51, 56, 72, 80, 100, 120, 140, 160, 180, 200, 500 and 1000. For each of the 22 × 62 = 1364 combinations of sequencing reads and cells sorted we generated 50 Flow-seq replicates to account for sampling variation.

### Deep learning architectures

Different neural networks architectures were implemented using PyTorch [47] and were systematically compared through a medium hyper-parameter optimisation (1000 examples using the Optuna default protocol [48], full details available in rebeca/hpo.py). The following architectures were considered: a multi-layer perceptron (MLP), a Convolutional Neural Network with an MLP used as a feature extractor in the final step, a Long Short Term Memory Recurrent Neural Network (RNN) and finally a hybrid CNN+RNN. The best performing model was a hybrid CNN+RNN with the following architecture: 2 layers of 1D CNN+maxpooling (256 channels, CNN kernel of size 4, max pooling kernel of size 2), followed by one biLSTM layer (hidden size 200). The output of the BiLSTM layer was flattened and followed by 2 fully-connected feed-forward layers with 512 units before the last layer comprising 8 units to predict the sequencing mode bin. Code for implementing architectures and model checkpoints are available in rebeca/model/sequence model.py and rebeca/checkpoints/convbilstm (**Data Availability**)

### Training and evaluation

A mini batch of 32 examples was used during training to minimize the crossentropy loss using the Adam optimizer with a learning rate of 3 × 10^−4^. Early stopping was used to identify over-fitting for each model. Training was conducted using two NVIDIA RTX 6000 GPUs. Evaluation was carried out by computing accuracy, precision, recall, F1 score and the confusion matrix. For each sequence subjected to data augmentation, we selected during testing the most frequent target output.

### General computational tools

All analysis scripts and the FORECAST package were written in Python version 3.9, using the numpy version 1.19.15 [49] for matrix algebra, scipy version 1.4.1 for statistical functions [50], numdifftools version 0.9.40 [51] for computing finite differences, joblib version 1.1.0 for multiprocessing, Pytorch version 1.12 [47] for deep learning. Statistical inference speed comparisons between MOM and ML approaches were conducted on an Apple iMac computer with a 3 GHz 6-core Intel Core i5 processor and 16 GB of RAM.

## Data Availability

Code for the FORECAST Python package is available at https://gitlab.com/Pierre-Aurelien/forecast. Code for modelling the deep neural networks is available at https://gitlab.com/Pierre-Aurelien/rebeca.

## Acknowledgments

This work was supported by the EPSRC/BBSRC Centre for Doctoral Training in Synthetic Biology grant EP/L016494/1 (P.-A.G.), BrisEngBio, a UKRI-funded Engineering Biology Research Centre grant BB/W013959/1 (T.E.G.), UKRI grant BB/W012448/1 (T.E.G.), a Turing Fellowship from The Alan Turing Institute under EPSRC grant EP/N510129/1 (T.E.G.), and a Royal Society University Research Fellowship grant UF160357 (T.E.G.)

## Author Contributions

P.-A.G. developed the methodology, carried out all computational work and performed the analysis. T.E.G. supervised the work. Both authors conceived of the study and wrote the manuscript.

