## Supplementary information for "Effective design and inference for cell sorting and sequencing based massively parallel reporter assays"

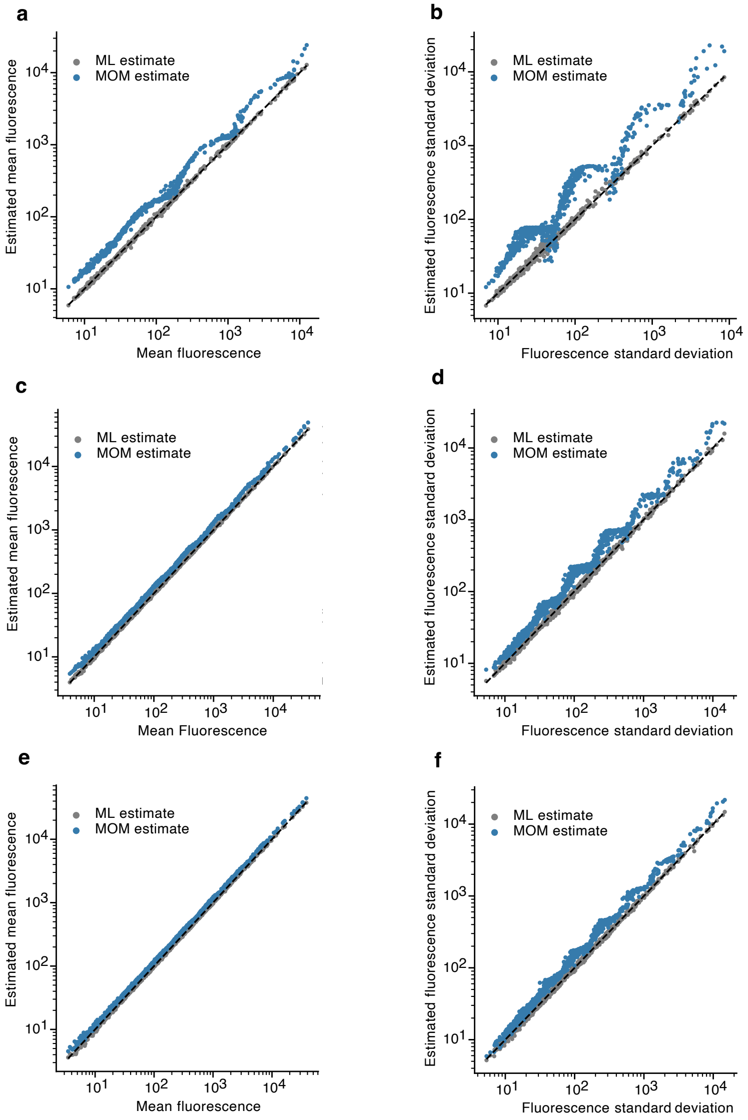

**Supplementary Figure 1: A staircase effect results when a few sorting bins are used.** Estimates for the fluorescence mean and standard deviation are compared with the ground truth when using an increasing number of sorting bins. Each point corresponds to a genetic variant colored blue for the Method of Moments (MOM) estimator, and grey for the Maximum Likelihood (ML) estimator. The Black dashed line shows $y = x$ (i.e., perfect estimation). (**a, b**) Cells are sorted into 6 bins (**c, d**) Cells are sorted into 8 bins. (**e, f**) Cells are sorted into 12 bins.The left scatter plots show the actual and estimated fluorescence mean for each genetic variant. The right scatter plots show the actual and estimated fluorescence standard deviation for each genetic variant.

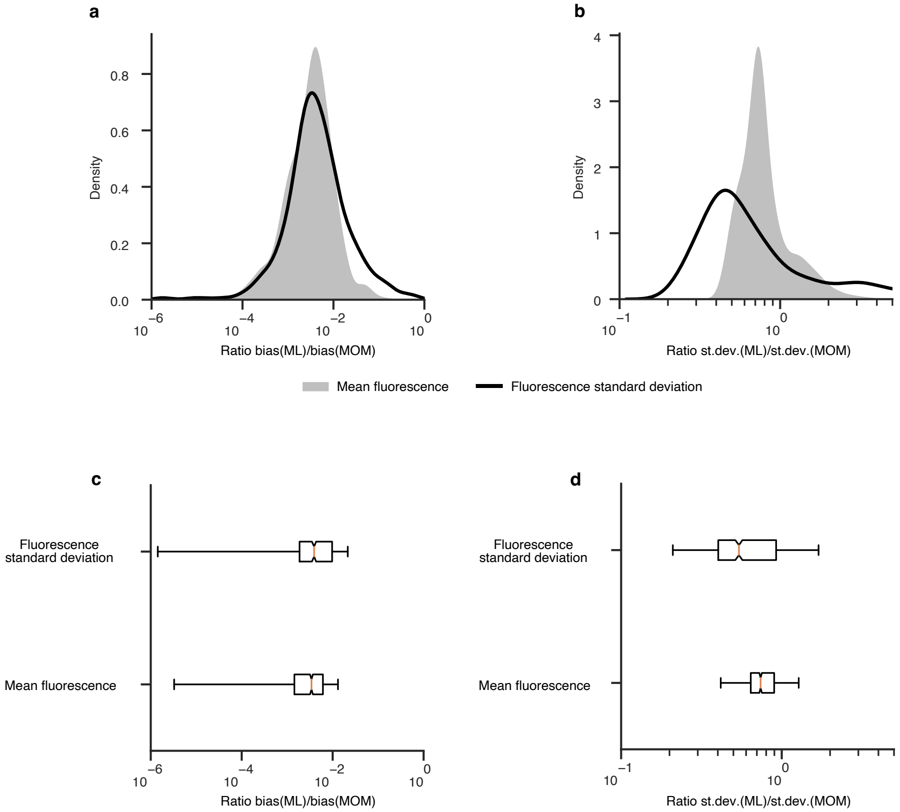

**Supplementary Figure 2: Sampling variation between the MOM and ML inference methods.** Sampling variation of MOM and ML inference was estimated by carrying out 500 synthetic Flow-seq simulations using biologically realistic expression characteristics (**Methods**). (**a**) Kernel density estimation of the ratio between the bias of the ML estimator and the bias of the MOM estimator when estimating the fluorescence mean or standard deviation across simulations. (**b**) Kernel density estimation of the ratio between the standard deviation of the ML estimator and the standard deviation of the MOM estimator when estimating the fluorescence mean or standard deviation across simulations. (**c**) Boxplot summarising the data shown in panel a. (**d**) Boxplot summarising the data in panel c.

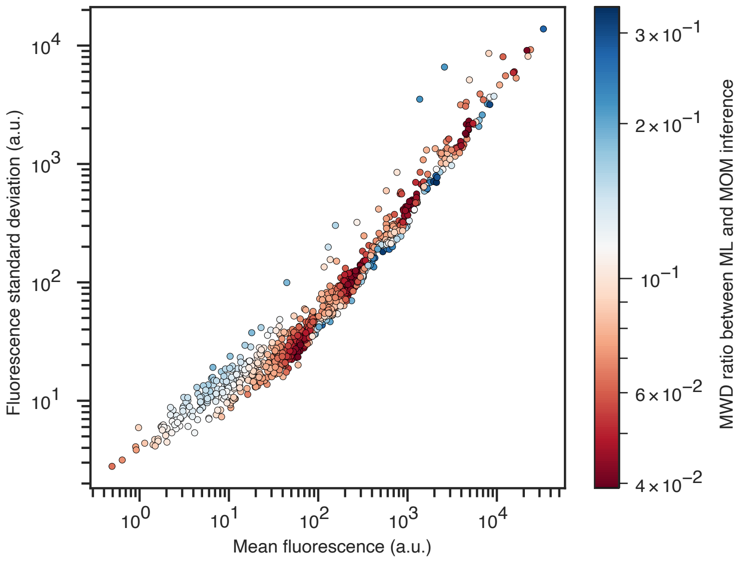

**Supplementary Figure 3: Comparing estimators.** ML estimator dominates the MoM estimator. Each circle represents a genetic construct, colored according to the Mean Wasserstein Distance (MWD) ratio between ML and MoM inference. The fluorescence mean and fluorescence standard deviation of each construct were used as coordinates for plotting this ratio. In all cases, this ratio is smaller than 1 (maximum at 0.34), indicating the superior performance of the ML method compared to the MoM method. Each genetic variant was characterized using a simulated Flow-seq experiment with 1018 genetic variants, $10^7$ sequencing reads, $10^6$ cells sorted, $10^2$ PCR amplification factor, 8 bins for cell sorting, and maximum fluorescence $f_{max} = 10^5$ using the fluorescence distribution parameters from the Taniguchi library (**Methods**). 500 different replicate simulations were generated to compute the mean 1-Wassertein distance between the inferred and ground-truth distribution when using ML and MOM inference (**Methods**).

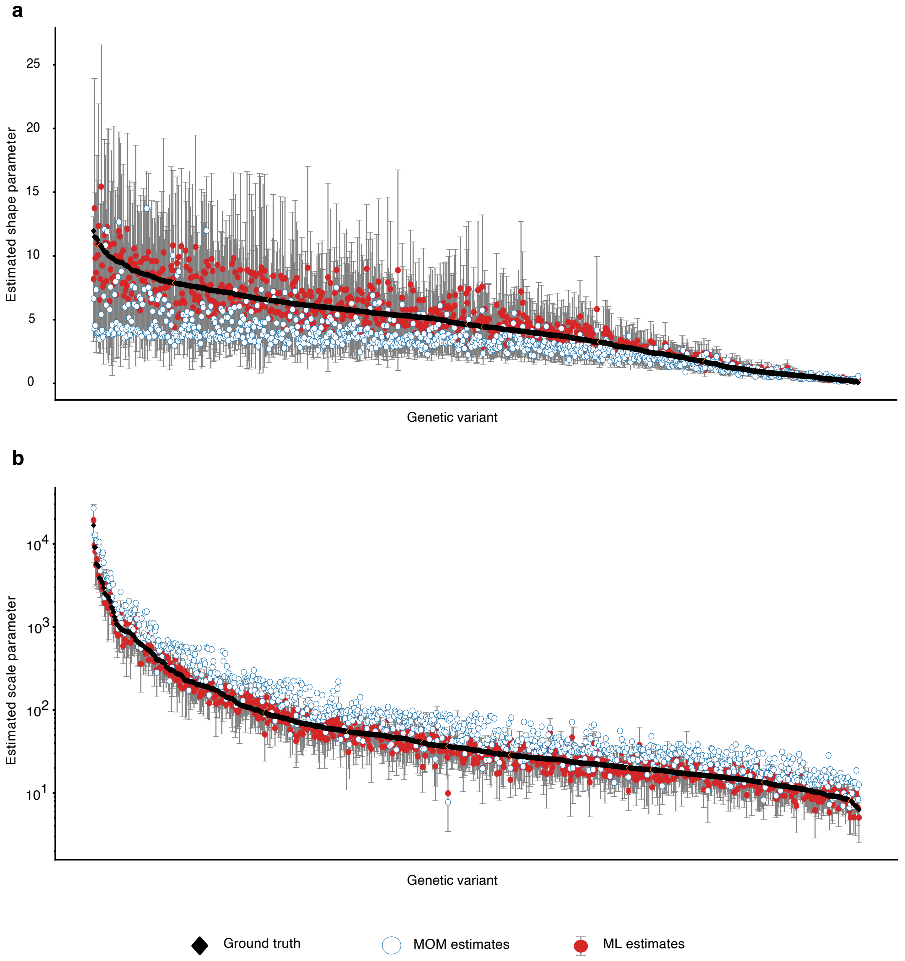

**Supplementary Figure 4: Comparing the MOM and ML estimates across an entire genetic library.** Estimated shape parameter (**a**) and estimated scale parameter (**b**) for the Gamma distribution of each genetic variant. Each genetic variant was characterized using a simulated Flow-seq experiment with 1018 genetic variants, $10^7$ sequencing reads, $10^6$ cells sorted, $10^2$ PCR amplification factor, 8 bins for cell sorting, and maximum fluorescence ($f_{max} = 10^5$) using the fluorescence distribution parameters from the Taniguchi library (**Methods**). Points show the inferred parameter value for the MOM (blue outline, white centre) and ML (red filled) methods, with the ground truth shown by a black-filled diamond. For the ML values, error bars show the 95% confidence interval for the estimate.

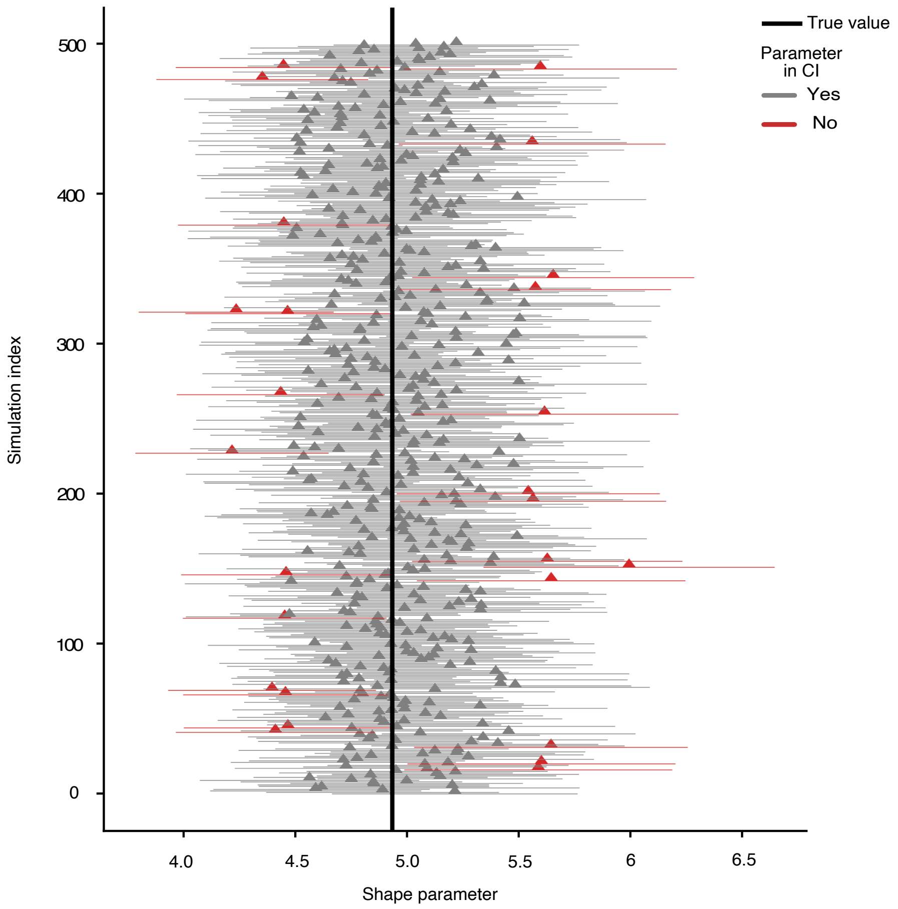

**Supplementary Figure 5: Coverage rate for the ML inference method.** Estimates of the fluorescence shape parameter for one genetic construct are plotted with their associated 95% confidence interval. The genetic construct was characterized after simulating a Flow-seq experiment with 1018 genetic variants, $10^7$ sequencing reads,$10^6$ cells sorted, $10^2$ PCR amplification factor, 8 bins for sorting cells, and maximum fluorescence ($f_{max} = 10^5$) using the fluorescence distribution parameters from the Taniguchi library (**Methods**). 500 different replicate simulations were generated to compute the coverage rate of the confidence interval, which as expected is also 95%.

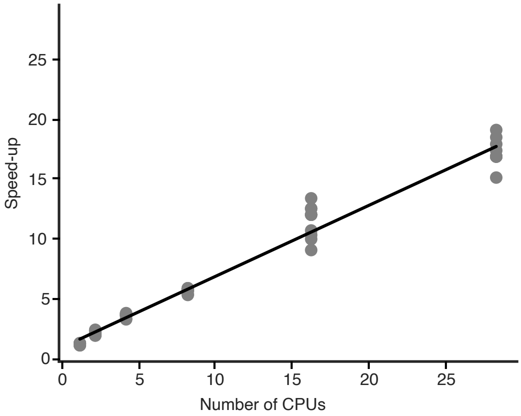

**Supplementary Figure 6: Scalability of the parallelized ML estimator.** Measuring scaling performance for ML inference to increasing numbers of CPUs. Each point corresponds to an independent measurement. The fitted observed linear speedup (solid black line) has a gradient of 0.6.

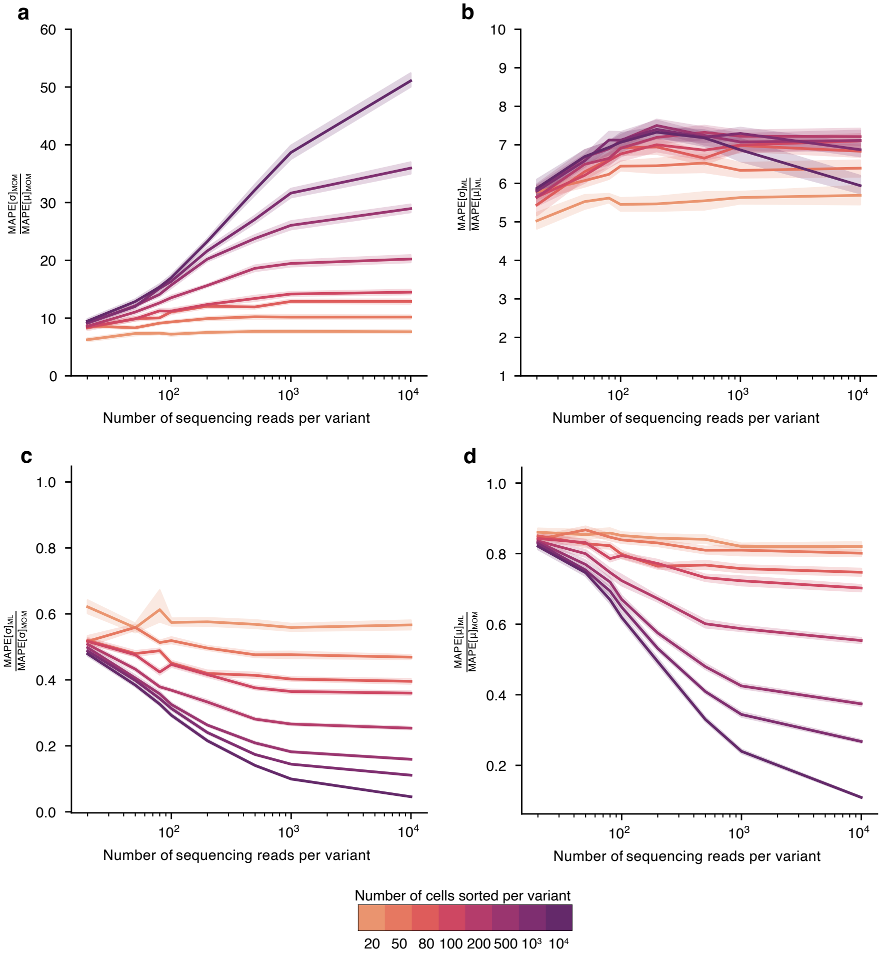

**Supplementary Figure 7: Differences in the accuracy of inferring fluorescence mean and standard deviation.** Fluorescence standard deviation estimation is less accurate then fluorescence mean estimation: The ratio between the MAPE of the fluorescence standard deviation and the MAPE of the fluorescence mean when using (**a**) MOM inference or (**b**) ML inference is always greater than 1. Inference using ML always yields a smaller relative error than inference using MOM with the ratio between the MAPE using ML and MAPE using MOM always being smaller than 1 when estimating the (**c**) fluorescence standard deviation and (**d**) the fluorescence mean.

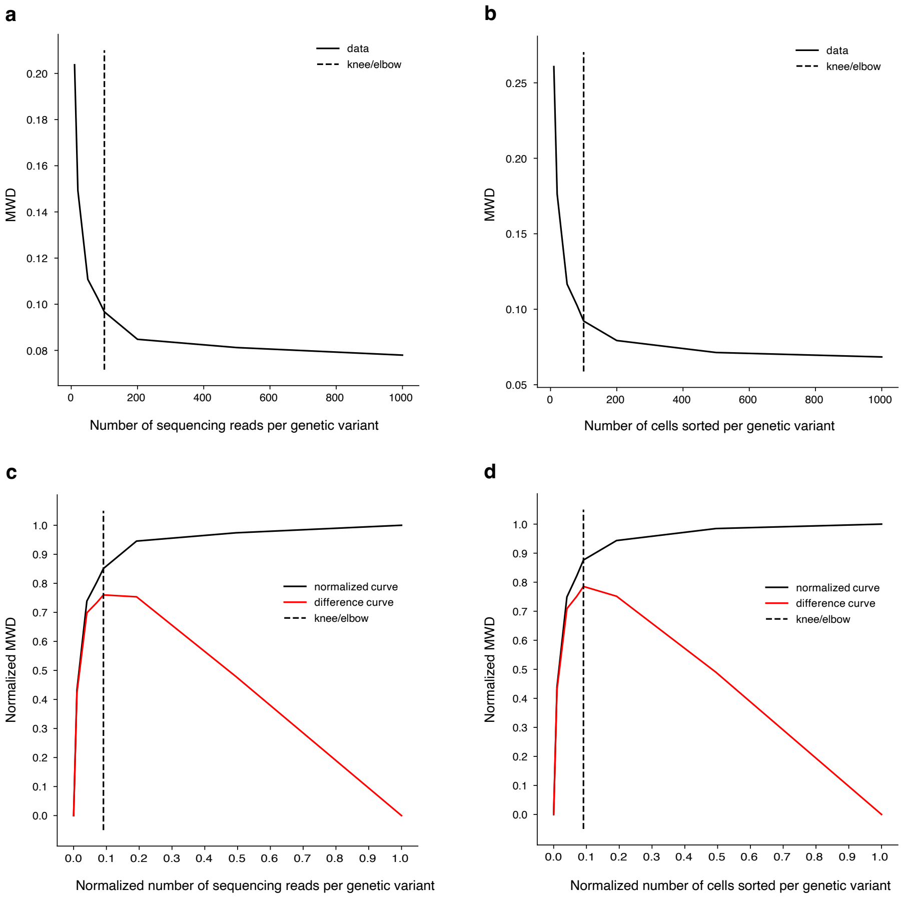

**Supplementary Figure 8: Finding optimal number of cells and sequencing reads.** (**a**) Mean 1-Wasserstein distance (MWD) between MOM-inferred and ground truth fluorescence distributions as a function of the number of sequencing reads. We picked the simulation with 80 cells per genetic variant during the cell sorting. (**b**) Mean 1-Wasserstein distance (MWD) between MOM-inferred and ground truth fluorescence distributions as a function of the number of cells sorted.We picked the simulation with 80 sequencing reads per genetic variant during the sequencing step. (**c**) Normalised version of panel a. (**d**) Normalised version of panel c. The Mean 1-Wasserstein distance was obtained by averaging over all genetic constructs in the Taniguchi library and all Flow-seq replicates (**Methods**).

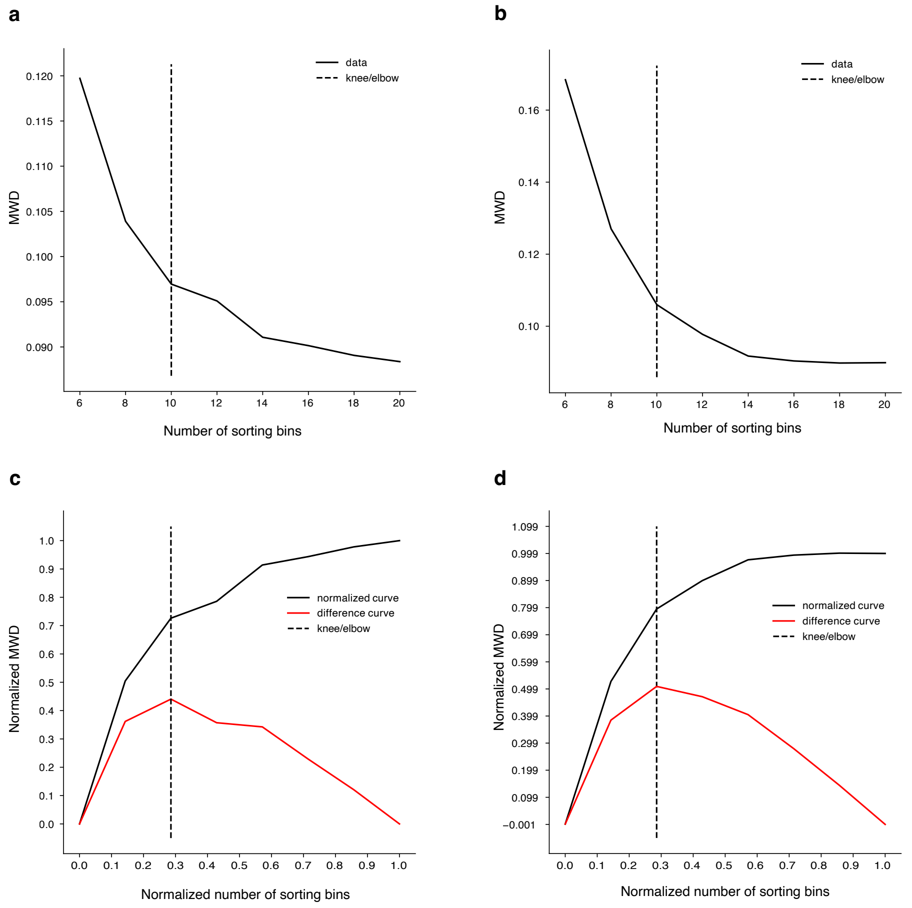

**Supplementary Figure 9: Finding optimal number of bins for cell sorting.** (**a**) Mean 1-Wasserstein distance (MWD) between ML-inferred and ground truth fluorescence distributions as a function of the number of sorting bins. (**b**) Mean Wasserstein distance between MOM-inferred and ground truth fluorescence distribution as a function of the number of sorting bins. (**c**) Normalised version of panel a. (**d**) Normalised version of panel b. The Mean 1-Wasserstein distance was obtained by averaging over all genetic constructs in the Taniguchi library and all Flow-seq replicates. We performed the simulation with the optimal number of sequencing reads (100 reads per genetic variant) and sorted cells (100 cells per genetic variant).

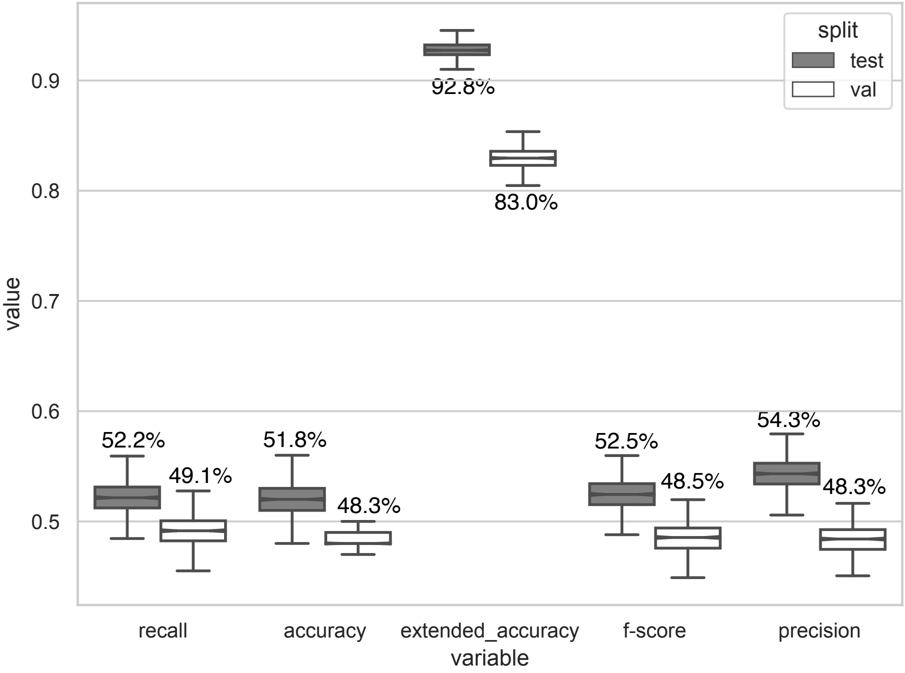

**Supplementary Figure 10: Evaluating the hybrid CNN+RNN neural network performance.** Performance across key metrics on the validation (white) and test (grey) datasets. Extended accuracy refers to the accuracy of the mode bin prediction allowing for a single bin deviation (i.e., allowing for incorrect predictions if they fall in a neighbouring bin). 95% confidence intervals were computed using 1000 bootstrap samples from each dataset.
